# Differential response to prey quorum signals indicates predatory range of myxobacteria

**DOI:** 10.1101/2021.06.04.447097

**Authors:** Shukria Akbar, Sandeep K. Misra, Joshua S. Sharp, D. Cole Stevens

## Abstract

A potential keystone taxa, myxobacteria contribute to the microbial food web as generalist predators. However, the extent of myxobacterial impact on microbial community structure remains unknown. The chemical ecology of these predator-prey interactions provides insight into myxobacterial production of biologically active specialized metabolites used to benefit consumption of prey as well as the perception of quorum signals secreted by prey. Using comparative transcriptomics and metabolomics, we compared how the predatory myxobacteria *Myxococcus xanthus* and *Cystobacter ferrugineus* respond to structurally distinct exogenous quorum signaling molecules. Investigating acylhomoserine lactone (AHL) and quinolone type quorum signals used by the clinical pathogen *Pseudomonas aeruginosa*, we identified a general response to AHL signals from both myxobacteria as well as a unique response from *C. ferrugineus* when exposed to the quinolone signal 4-hydroxy-2-heptylquinolone (HHQ). Oxidative detoxification of HHQ in *C. ferrugineus* was not observed from *M. xanthus*. Subsequent predation assays indicated *P. aeruginosa* to be more susceptible to *C. ferrugineus* predation. These data indicate that as generalist predators myxobacteria demonstrate a common response to the ubiquitous AHL quorum signal class, and we suggest this response likely involves recognition of the homoserine lactone moiety of AHLs. We also suggest that oxidation of HHQ and superior predation of *P. aeruginosa* observed from *C. ferrugineus* provides an example of how prey signaling molecules impact predatory specialization of myxobacteria by influencing prey range.

**Summary:** Multiomic analysis of transcriptional and metabolic responses from the predatory myxobacteria *Myxococcus xanthus* and *Cystobacter ferrugineus* exposed to prey signaling molecules of the acylhomoserine lactone and quinolone quorum signaling classes provided insight into myxobacterial specialization associated with predatory eavesdropping. We suggest that the general response observed from both myxobacteria exposed to acylhomoserine lactone quorum signals is likely due to the generalist predator lifestyles of myxobacteria and ubiquity of acylhomoserine lactone signals. We also provide data that indicates the core homoserine lactone moiety included in all acylhomoserine lactone scaffolds to be sufficient to induce this general response. Comparing both myxobacteria, unique transcriptional and metabolic responses were observed from *Cystobacter ferrugineus* exposed to the quinolone signal 4-hydroxy-2-heptylquinoline (HHQ) natively produced by *Pseudomonas aeruginosa*. We suggest that this unique response and ability to metabolize quinolone signals contribute to the superior predation of *P. aeruginosa* observed from *C. ferrugineus*. These results further demonstrate myxobacterial eavesdropping on prey signaling molecules and provide insight into how responses to exogenous signals might correlate with prey range of myxobacteria.

**Originality-Significance Statement:** This manuscript provides the first multiomic analysis of how predatory myxobacteria respond to exogenous prey signaling molecules and details the differences observed by comparing responses from two myxobacteria.

## Introduction

The uniquely multicellular lifestyles of myxobacteria have motivated continued efforts to explore the myxobacterium *Myxococcus xanthus* as a model organism for cooperative behaviors including development (Islam et al., 2020; Sharma et al., 2021), motility (Mercier et al., 2020; Rendueles and Velicer, 2020; Zhang et al., 2020b), and predation (Thiery and Kaimer, 2020; Zhang et al., 2020a; Sydney et al., 2021). Often attributed to their need to acquire nutrients as generalist predators (Nair et al., 2019) and capacity to prey upon clinical pathogens (Livingstone et al., 2017), myxobacteria have also been a valuable resource for the discovery of novel specialized metabolites as potential therapeutic lead compounds (Herrmann et al., 2017; Baltz, 2019; Perez et al., 2020). The diversities in structural scaffolds and observed activities as well as the unique chemical space associated with myxobacterial metabolites when compared with more thoroughly explored Actinobacteria make myxobacteria excellent sources for efforts focused on the discovery of therapeutics (Herrmann et al., 2017; Baltz, 2019). However, the connection between myxobacterial predation and production of these biologically active metabolites remains underexplored. Currently, only the metabolites myxovirescin (Xiao et al., 2011; Ellis et al., 2019; Wang et al., 2019) and myxoprincomide (Cortina et al., 2012; Muller et al., 2016) have been directly implicated to be involved during *Myxococcus xanthus* predation of *Escherichia coli* and *Bacillus subtilis* respectively. In fact, the chemical ecology of predator-prey interactions between myxobacteria and prey remains underexplored (Findlay, 2016). The predatory capacity or prey range of myxobacteria cannot be directly correlated with phylogeny (Livingstone et al., 2017; Arend et al., 2020). Presently, the best determinants for broadly assessing prey ranges are genetic features that might provide specific traits to overcome predation resistances of individual prey. For example, myxobacteria possessing the formaldehyde dismutase gene *fdm* demonstrated comparatively better predation of toxic formaldehyde secreting *Pseudomonas aeruginosa* a clinical pathogen observed to be somewhat recalcitrant to myxobacterial predation (Arend et al., 2020).

The recent observation that acylhomoserine lactone (AHL) quorum signals from prey microbes impact the predatory capacity of *M. xanthus* suggests that quorum signals might influence predator-prey interactions (Lloyd and Whitworth, 2017). Although two orphaned, functional AHL synthases have been reported, no myxobacteria have been observed to produce AHLs (Albataineh et al., 2021). However, a recent survey of signaling systems within the family Myxococcaceae reported the presence of conserved AHL receptor (LuxR) homologs and inferred that many myxobacteria within the 2 genera *Myxococcus* and *Corallococcus* are capable of sensing AHL signaling molecules (Whitworth and Zwarycz, 2020). While this suggests that predatory myxobacteria might eavesdrop on prey quorum signaling, the observed reaction from *M. xanthus* might also simply be the result of exogenous AHLs as a nutritional gradient. Herein we utilize a combination of transcriptomics and metabolomics to determine how myxobacterial responses to quorum signals produced by *P. aeruginosa* might indicate predatory capacity.

By exposing myxobacteria to structurally and functionally distinct classes of prey quorum signals comparing ubiquitous AHL signals and quinolone signals more unique to pseudomonads (Papenfort and Bassler, 2016), we anticipated that a differential response exclusive to a specific signal class would support predatory eavesdropping and perhaps correlate with improved predation of *P. aeruginosa*. For these experiments we exposed each myxobacterium to AHL signals (Galloway et al., 2011) as well as the quinolone signal 4-hydroxy-2-heptylquinolone (HHQ) (Deziel et al., 2004; Dubern and Diggle, 2008; Garcia-Reyes et al., 2020). Ubiquitous to Proteobacteria (notably excluding myxobacteria) and numerous other non-Proteobacteria genera, AHLs are the most common class of quorum signal autoinducers and are often implicated in interspecies communication within polymicrobial communities (Shiner et al., 2005; Mukherjee and Bassler, 2019). Also associated with the modulation of interspecies and interkingdom behaviors (Reen et al., 2011), the quinolone signal HHQ contributes to the pathogenicity of *P. aeruginosa* by participating in the regulation of various virulence factors (Dubern and Diggle, 2008; Reen et al., 2011). Exploration of the myxobacterial response to prey quorum signals not only provides insight into the impact of shared chemical signals might have on predator-prey interactions within bacterial communities but may also provide further genetic determinants that indicate predatory capacities of myxobacteria.

As a model organism for developmental studies, *M. xanthus* is the best characterized myxobacterium and has already demonstrated a behavioral response to exogenous AHLs (Lloyd and Whitworth, 2017). However, we suspected that routine use as a laboratory strain, constitutive toxicity (Livingstone et al., 2018), and well-explored specialized metabolism (Cortina et al., 2012; Herrmann et al., 2017) of *M. xanthus* might diminish its viability as the sole myxobacterium for these experiments. Therefore, *Cystobacter ferrugineus* was also included as a more recent myxobacterial isolate with a less explored biosynthetic capacity and prey range (Akbar et al., 2017; Goes et al., 2020). Both *M. xanthus* and *C. ferrugineus* have an annotated solo LuxR-type AHL receptor present in their genomes (WP_011555271.1 and WP_071900454.1) (Subramoni and Venturi, 2009; Tobias et al., 2020; Xu, 2020). However, homology-based annotation of these features only indicates the helix-turn-helix DNA-binding domain of LuxR receptors, and neither include an AHL-binding site motif (PF03472) (Baikalov et al., 1996; Vannini et al., 2002; Mukherjee and Bassler, 2019). Despite the absence of a canonical receptor, exogenous AHLs have been observed to stimulate the motility and predatory activity of *M. xanthus* (Lloyd and Whitworth, 2017). Also of note, neither *M. xanthus* or *C. ferrugineus* possess a homologous PqsR-type HHQ receptor (Diggle et al., 2003; Wade et al., 2005). Utilizing a multiomic approach to assess the transcriptomic and metabolomic responses of *M. xanthus* and *C. ferrugineus* when exposed to AHL and quinolone signals, we sought to determine if structurally and functionally dissimilar quorum signals from prey elicit distinct responses from predatory myxobacteria that correlate with successful predation of *P. aeruginosa*.

## Results

### C6-AHL induces a general transcriptional response from both *M. xanthus* and *C. ferrugineus*

Exposure experiments utilizing a concentration of C6-AHL previously shown to elicit a predatory response from *M. xanthus* (9 μM) (Lloyd and Whitworth, 2017) were conducted in triplicate for both *M. xanthus* and *C. ferrugineus* with DMSO exposures serving as vehicle, negative controls for comparative analysis. Comparative transcriptomic analysis from RNAseq data revealed C6-AHL exposure impacted transcription of a total of 76 genes from *C. ferrugineus* experiments and just nine genes from *M. xanthus* experiments when only considering a ≥4-fold change in transcription at p ≤0.01 (Figure 1, Supplemental Dataset 1). For this reason, our analysis of *M. xanthus* exposure experiments was expanded to include significant features at p ≤0.05 resulting in an updated total of 59 impacted features from C6-AHL exposure (Figure 1). While this indicates less variability across *C. ferrugineus* exposure experiments, we contend that this expansion provides a broader and more thorough analysis of statistically significant impacted features for our analysis. A similar consideration of *C. ferrugineus* genes with ≥4-fold change in transcription at p ≤0.05 by C6-AHL exposure would provide an additional 119 impacted genes for consideration (Supplemental Dataset 1). *M. xanthus* features included at the more stringent significance cutoff of p ≤0.01 are indicated in Figure 1. These data revealed that C6-AHL exposure elicited a general downregulation of genes with a total of 55 downregulated genes observed from *M. xanthus* and 51 genes from *C. ferrugineus*. Only one gene was observed to be upregulated by *M. xanthus* when exposed to C6-AHL, and 25 total upregulated genes were observed from *C. ferrugineus* during C6-AHL exposure.

**Figure 1:**
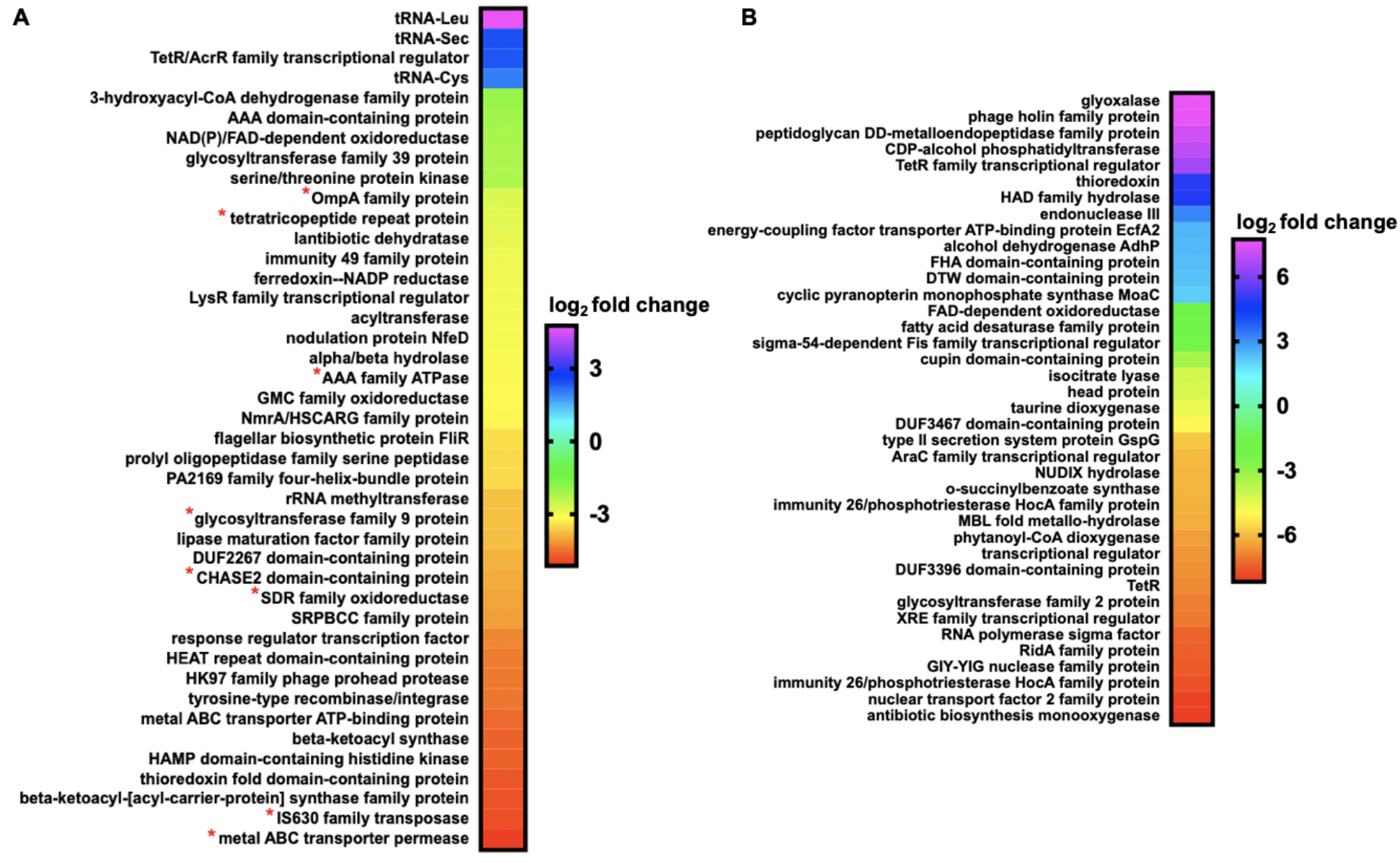
Transcriptomic data from myxobacteria exposed to C6-AHL. (A) Differentially expressed genes and features from *M. xanthus* exposed to C6-AHL when compared to signal unexposed *M. xanthus* control (p ≤0.05); * indicates features also impacted at p ≤0.01. (B) Differentially expressed genes from *C. ferrugineus* exposed to C6-AHL when compared to signal unexposed *C. ferrugineus* control (p ≤0.01). Data depicted as an average log2 fold change from three biological replicates. Impacted features annotated as hypothetical not included.

Comparing annotated features impacted by C6-AHL exposure across both datasets and their putative roles by general system, numerous features involved in signal transduction pathways and transcriptional regulation were included in both datasets with seven regulatory features downregulated by *M. xanthus* and six downregulated by *C. ferrugineus* (Figure 2). Both myxobacteria also had a TetR family transcriptional regulator upregulated by C6-AHL exposure. Multiple features associated with primary and specialized metabolisms and cell wall biogenesis and maintenance were downregulated by C6-AHL exposure across both datasets. Considering previous reports that C6-AHL exposure suppresses *M. xanthus* sporulation (Lloyd and Whitworth, 2017), we sought to determine if C6-AHL exposure effected either of the transcriptional regulators associated with *M. xanthus* sporulation FruA or MrpC (Ogawa et al., 1996; Robinson et al., 2014; Marcos-Torres et al., 2020). While no significant change in FurA was observed, transcription of the gene product MrpC was downregulated 1.7-fold by *M. xanthus* exposure to C6-AHL. However, transcription of the FruA (WP_071904077.1) or MrpC (WP_071900118) homologs from *C. ferrugineus* was not significantly changed by C6-AHL exposure. While no obvious predatory features associated with motility or lytic enzymes were directly impacted in our C6-AHL exposed *M. xanthus* results, we suspect that this could be due to the previously reported constitutive toxicity of *M. xanthus* observed in both the presence and absence of prey (Livingstone et al., 2018). The increased transcription of lytic enzymes and mobile genetic elements observed from *C. ferrugineus* exposed to C6-AHL suggest a predatory response; however, these features could also be associated with a defense response akin to phage defense. Transcription of neither of the annotated LuxR-type receptors (*M. xanthus*, WP_011555271.1; *C. ferrugineus*, WP_071900454.1) was affected by C6-AHL exposure. Overall considering the most significantly impacted features across both datasets, C6-AHL exposure elicited somewhat similar responses from both myxobacteria including numerous features associated with transcriptional regulation and signal transduction, primary and specialized metabolisms, and cell wall maintenance.

**Figure 2:**
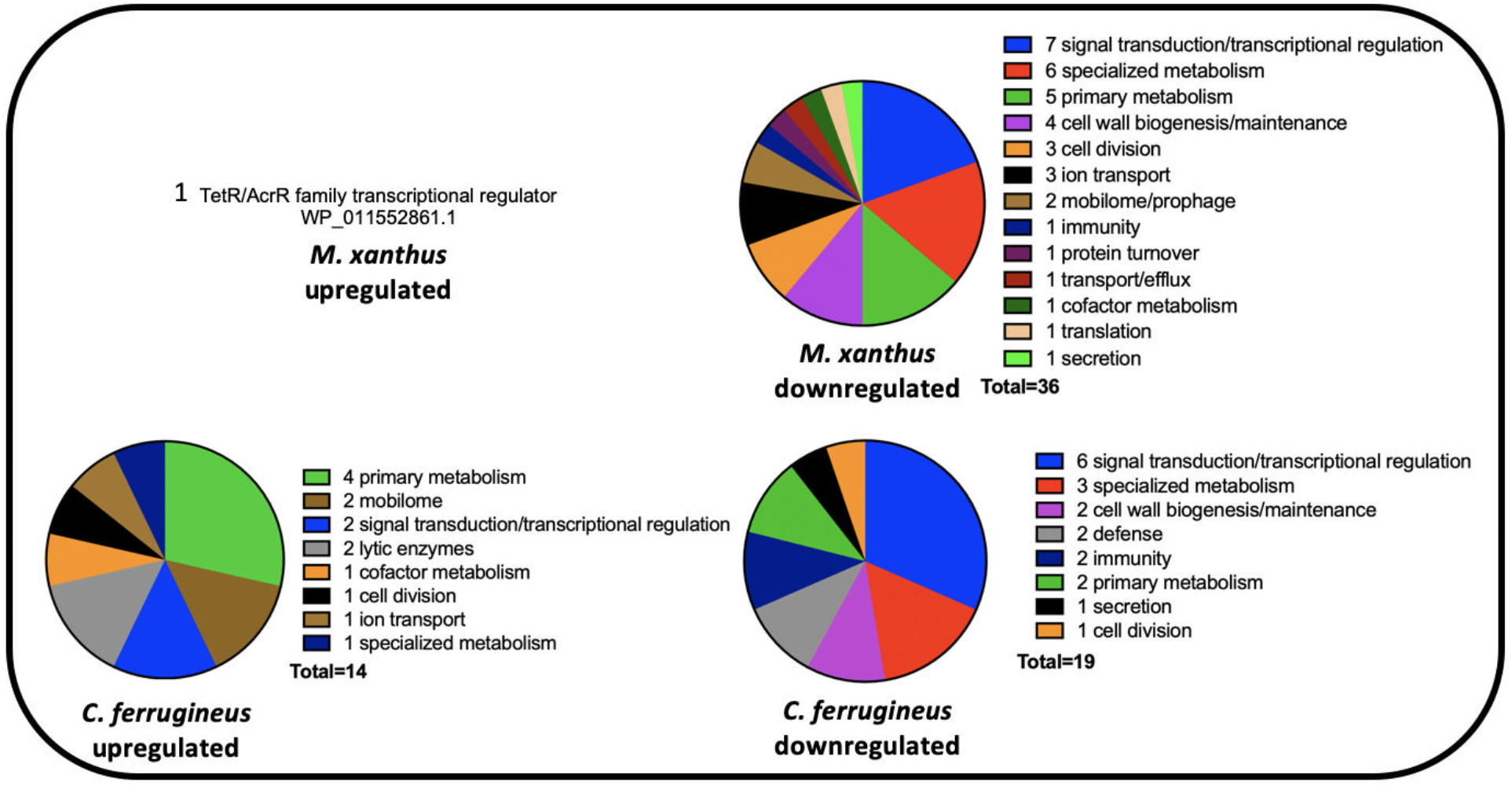
Putative roles of Prokaryotic Genome Annotation Pipeline (PGAP)-annotated genes impacted by C6-AHL exposure (from Figure 1) comparing *M. xanthus* and *C. ferrugineus*.

### HHQ elicits contrasting responses from *M. xanthus* and *C. ferrugineus*

Comparative transcriptomics from RNAseq data from exposure experiments with HHQ (9 μM) introduced to plates of *M. xanthus* and *C. ferrugineus* were also conducted in triplicate with DMSO exposures serving as HHQ unexposed, negative controls for comparative analysis. Comparative transcriptomic analysis from RNAseq data revealed HHQ exposure led to a ≥4-fold (p ≤0.05) change in transcription of a total of 186 genes from *C. ferrugineus* and 31 total genes from *M. xanthus* (Figure 3). Unlike the similar responses elicited by C6-AHL exposure, contrasting responses were apparent when comparing data between the myxobacteria. Data from *M. xanthus* experiments revealed overlap between responses to C6-AHL and HHQ with a total of nine upregulated genes and 22 downregulated genes including five genes also downregulated by C6-AHL exposure. Overlapping annotated features impacted by both C6-AHL and HHQ included a NmrA/HSCARG family protein (WP_011556972.1), an immunity 49 family protein (WP_011550233.1), a CHASE2 domain-containing protein (WP_011554259.1), and two hypothetical proteins (WP_011555268.1 and WP_011552217.1). Comparing impacted genes from AHL and HHQ exposure experiments, further overlap between putative roles of annotated genes was also observed from *M. xanthus* with multiple impacted genes predicted to be involved in transcriptional regulation and signal transduction and cell wall biogenesis and maintenance (Figure 2 and Figure 4). Of note, the pleiotropic regulator MrpC was also downregulated 2.2-fold in *M. xanthus* exposed to HHQ which is comparable to 1.7-fold downregulation of MrpC observed with C6-AHL exposure.

**Figure 3:**
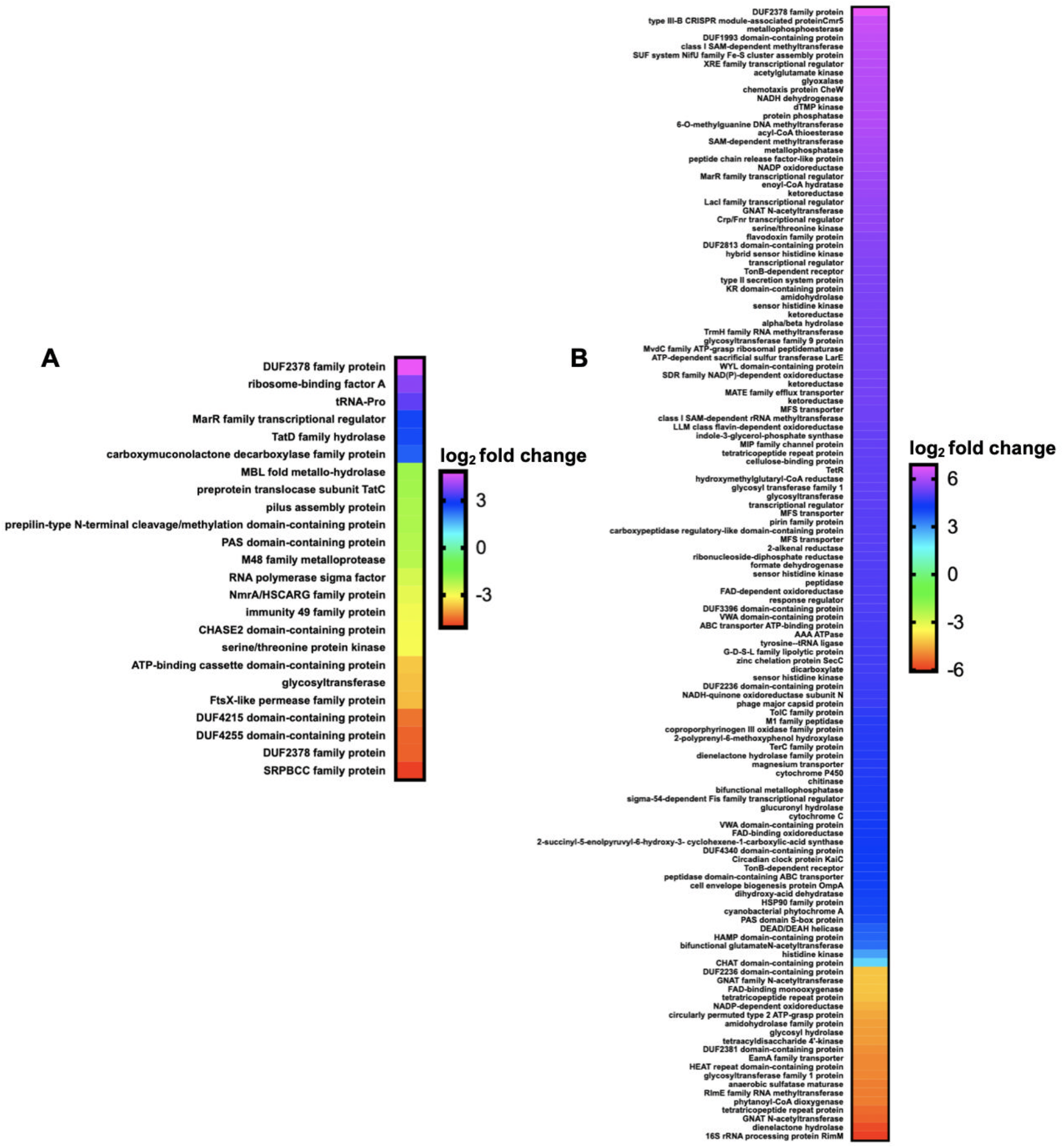
Transcriptomic data from myxobacteria exposed to HHQ. (A) Differentially expressed genes and features from *M. xanthus* exposed to HHQ when compared to signal unexposed *M. xanthus* control (p ≤0.05). (B) Differentially expressed genes from *C. ferrugineus* exposed to HHQ when compared to signal unexposed *C. ferrugineus* control (p ≤0.05). Data depicted as an average log2 fold change from three biological replicates. Impacted features annotated as hypothetical not included.

**Figure 4:**
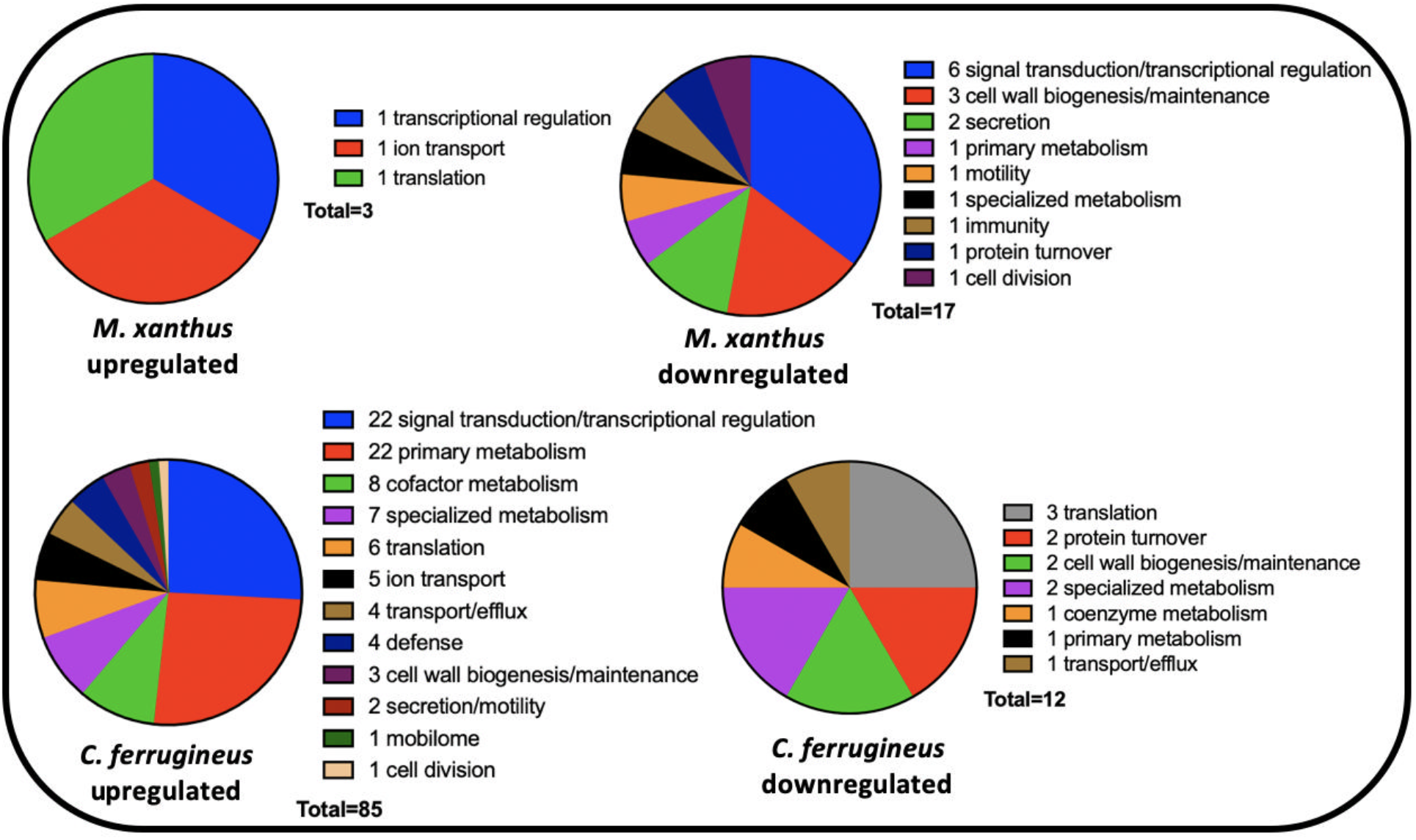
Putative roles of PGAP-annotated genes impacted by HHQ exposure (from Figure 3) comparing *M. xanthus* and *C. ferrugineus*.

Unlike the overlap in responses to both signals observed from *M. xanthus*, none of the 186 genes effected by HHQ exposure overlapped with the 76 genes impacted by C6-AHL exposure. Considering annotated genes by functional category, *C. ferrugineus* genes upregulated by HHQ exposure (156 total) were largely associated with signal transduction and transcriptional regulation, various metabolic pathways, and multiple genes associated with protein translation and turnover, cell wall biogenesis and maintenance, and specialized metabolism were downregulated (29 total) (Figure 4). Interestingly, an annotated FAD-dependent oxidoreductase (WP_071901324.1) homologous to the monooxygenase PsqH from *P. aeruginosa* (91% coverage; 38% identity) which hydroxylates HHQ to yield 2-heptyl-3,4-dihydroxyquinolone or pseudomonas quinolone signal (PQS) was upregulated 31-fold (Diggle et al., 2003; Ritzmann et al., 2021). An outlier to the contrasting responses to HHQ, an annotated DUF2378 family protein (*M. xanthus*, WP_011553830.1; *C. ferrugineus*, WP_084736518.1) was significantly upregulated in both myxobacteria; DUF2378 family proteins are ~200 amino acid proteins with no known function that are exclusive to myxobacteria. Overall, these results indicate that *M. xanthus* exhibits a similar transcriptional response to both C6-AHL and HHQ whereas HHQ elicits a distinct response from *C. ferrugineus* dissimilar from the more general response observed from both myxobacteria when exposed to C6-AHL.

### Differential metabolic impact of AHL and HHQ signals

Subsequent exposure experiments were conducted with *M. xanthus* and *C. ferrugineus* exactly as done for our RNAseq experiments with an additional AHL signal, 3-oxo-C6-AHL, also included. Crude, organic phase extracts generated from these experiments were subjected to untargeted mass spectrometry and the XCMS-MRM (v3.7.1) platform (Domingo-Almenara et al., 2018; Forsberg et al., 2018) was utilized for comparative analysis and determination of statistical significance for all detected features. Comparing features with significantly impacted intensities (p ≤0.02) during these signal exposure experiments, all three signals elicited a more apparent response from *C. ferrugineus* (Figure 5 and Supplemental Figure 1). Despite the comparatively diminished response from *M. xanthus*, two general trends were apparent when comparing the signals responses between both myxobacteria. First, C6-AHL and 3-oxo-C6-AHL exposure resulted in highly similar responses from both myxobacteria with few to no uniquely impacted features specific to either AHL signal (Figure 5). Second, HHQ exposure induced a dramatic change in the metabolic profile of *C. ferrugineus* that was not observed from HHQ-exposed *M. xanthus*. A total of 47 features from *C. ferrugineus* were impacted by both AHL signals while 133 features were affected by HHQ exposure. Intrigued by the difference in responses, additional experiments where both myxobacteria were exposed to exogenous C6-AHL and HHQ simultaneously were done. Comparative analysis of results revealed that the addition of C6-AHL did not dramatically impact the change in metabolic profile observed from either myxobacteria when exposed to HHQ (Figure 6). Conversely, impacted features observed in our previous AHL exposure experiments were not observed in our C6-AHL + HHQ experiments. For example, of the 47 total overlapping *C. ferrugineus* features with significantly changed intensities during AHL exposure conditions, 36 were not observed to change during C6-AHL + HHQ exposure experiments. From these results, we determined that both myxobacteria demonstrate a metabolic response unique to either HHQ or AHL-type chemical signals with a conserved response to both C6-AHL and 3-oxo-C6-AHL. These data also revealed a unique metabolic response from *C. ferrugineus* when exposed to HHQ similar to our previous transcriptomic observation.

**Figure 5:**
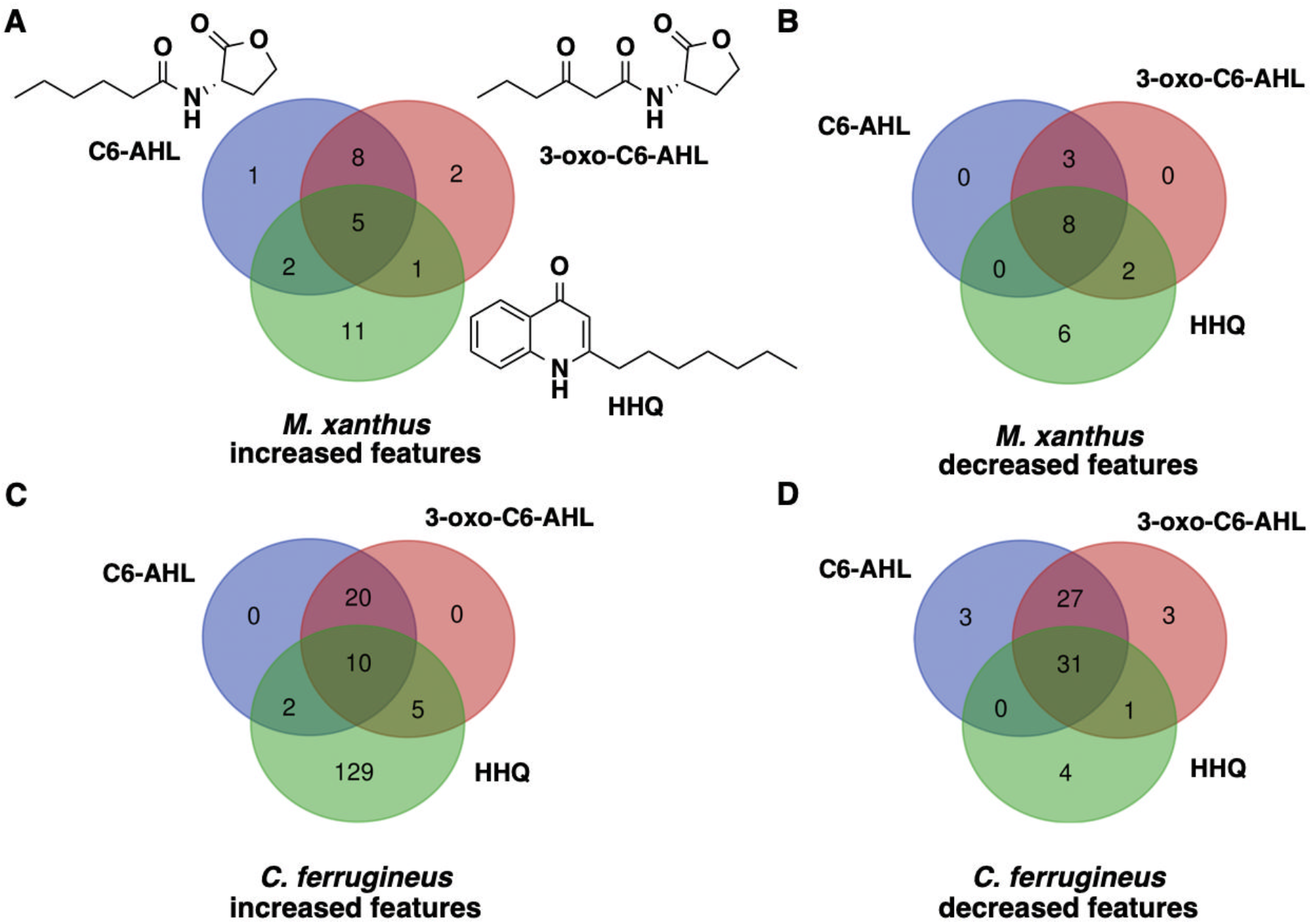
Comparison of metabolomic response to C6-AHL, 3-oxo-C6-AHL, and HHQ exposure experiments with *M. xanthus* (A and B) and *C. ferrugineus* (C and D). Numbers included in each Venn diagram account for a unique detected feature with a significantly impacted intensity upon exposure to the indicated signaling molecule provided by XCMS-multigroup analysis (n=3, p ≤0.02).

**Figure 6:**
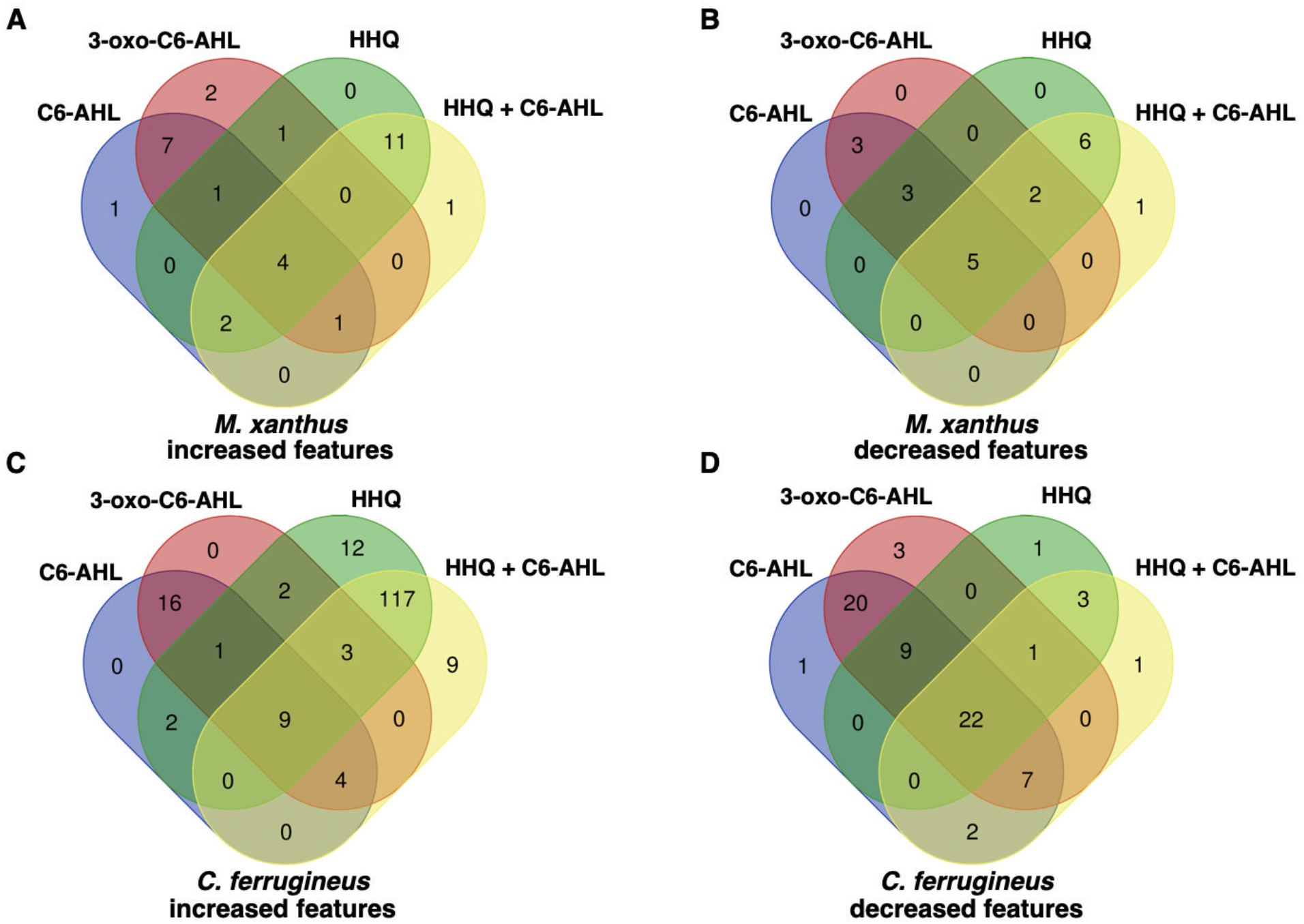
Comparison of metabolomic response to C6-AHL, 3-oxo-C6-AHL, and HHQ exposure experiments with *M. xanthus* (A and B) and *C. ferrugineus* (C and D) including additional C6-AHL + HHQ exposure experiments. Numbers included in each Venn diagram account for a unique detected feature with a significantly impacted intensity upon exposure to the indicated signaling molecule provided by XCMS-multigroup analysis (n=3, p ≤0.02).

### Conserved metabolomic response to AHLs and determination of core L-homoserine lactone elicitor

Intrigued by the overlap in metabolic responses observed from AHL signal exposure, we were curious if the core homoserine lactone moiety present in all AHL-type quorum signals was sufficient to elicit a similar response. Untargeted mass spectrometry and XCMS-MRM analysis of additional exposure experiments with *C. ferrugineus* including either L-homoserine lactone (L-HSL) the stereoisomer present in natural AHL-type quorum signals (Papenfort and Bassler, 2016; Mukherjee and Bassler, 2019), D-homoserine lactone (D-HSL) the enantiomer of L-HSL, or boiled C6-AHL were completed to determine any overlap with previously observed responses to C6-AHL and 3-oxo-C6-AHL exposure. Comparative analysis of statistically impacted features by signal intensity (p ≤0.02) revealed 20 overlapping features from the L-HSL, C6-AHL, and 3-oxo-C6-AHL exposure experiments and just three overlapping features from the L-HSL, D-HSL, C6-AHL, and 3-oxo-C6-AHL exposure experiments (Figure 7, Supplemental Figure 2). Hierarchical clustering of detected feature intensities from L-HSL, D-HSL, C6-AHL, and control datasets also revealed clustering between L-HSL and C6-AHL datasets (Supplemental Figure 3). These results suggest that the core homoserine lactone core present in all AHLs is sufficient for predatory eavesdropping by myxobacteria.

**Figure 7:**
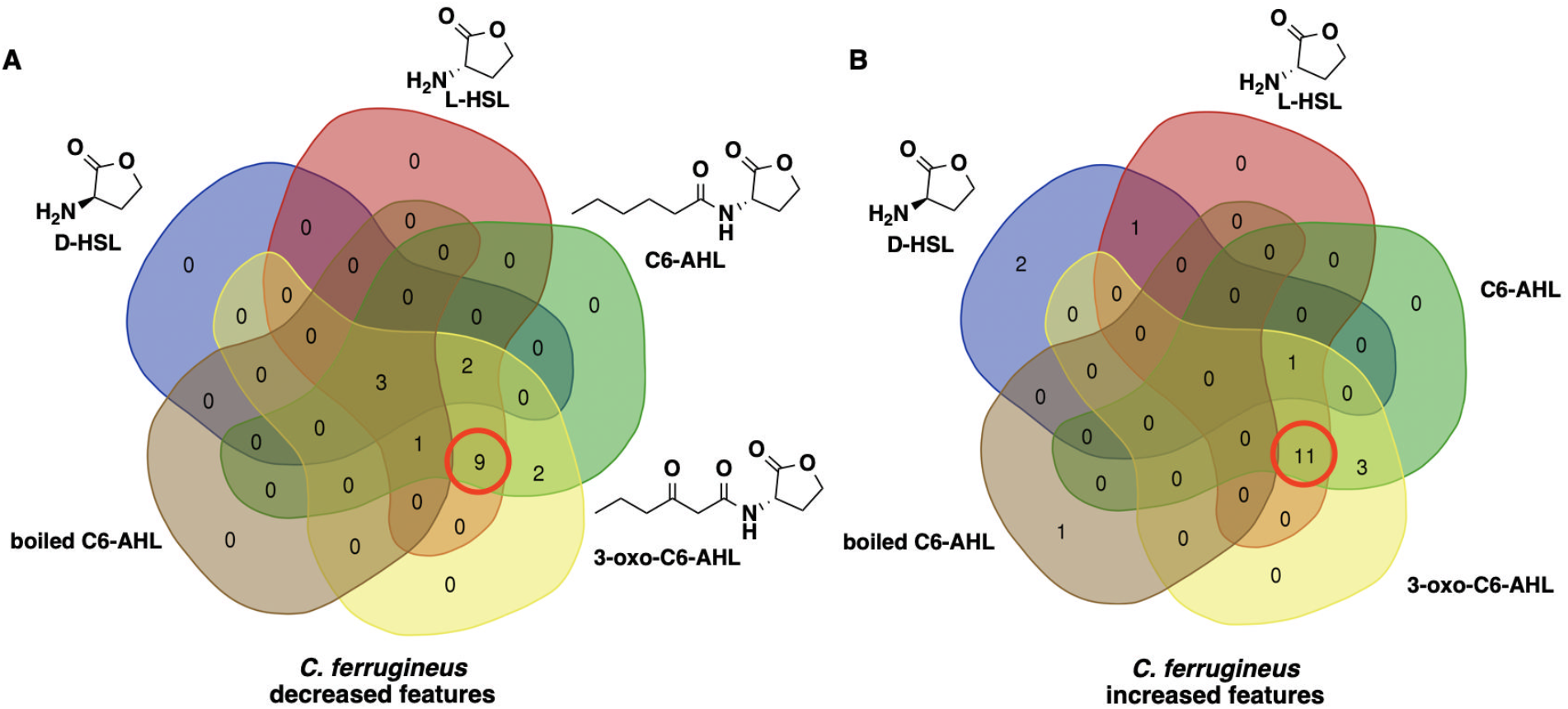
Overlap in metabolic response to C6-AHL, 3-oxo-C6-AHL, and L-HSL exposure observed from *C. ferrugineus*. Venn diagrams include the number of metabolic features with a significant increase (A) or decrease (B) in detected ion intensity compared to signal unexposed controls provided by XCMS-multigroup analysis (n=3; p ≤0.02). Red circles indicate overlapping metabolic features of L-HSL with C6-AHL and 3-oxo-C6-AHL.

### Oxidative detoxification of HHQ observed from *C. ferrugineus*

Comparing metabolomic datasets from HHQ exposure experiments, oxidized analogs of HHQ detected at 260.164 m/z were exclusive to the *C. ferrugineus* dataset. Authentic standards for the oxidized HHQ quinolone signals PQS and 2-heptyl-4-hydroxyquinoline *N*-oxide (HQNO) were used to determine that both oxidized signals were present in HHQ-exposed *C. ferrugineus* extracts (Figure 8, Supplemental Figure 4) (Cao et al., 2020). Oxidative detoxification of quinolone signals including HHQ has been reported from numerous bacteria (Thierbach et al., 2017; Ritzmann et al., 2021). Additional experiments exposing *C. ferrugineus* to either PQS or HQNO provided insight into a similar oxidative detoxification route for quinolone signals with HHQ observed to be oxidized to either HQNO or PQS. The presence of a metabolite with an exact mass and similar MS^2^ fragmentation pattern matching 2-heptyl-3,4-dihydroxyquinoline-*N*-oxide (PQS-NO) an oxidation product reported by Thierbach *et al*. was also observed in *C. ferrugineus* extracts from HHQ, PQS, and HQNO exposure experiments suggesting subsequent oxidation of both PQS and HQNO (Figure 8, Supplemental Figures 5 and 6) (Thierbach et al., 2017). These results suggest that *C. ferrugineus* possesses a detoxification route for quinolone signals not observed from *M. xanthus* and oxidizes the quinolone signals HHQ, PQS, and HQNO.

**Figure 8:**
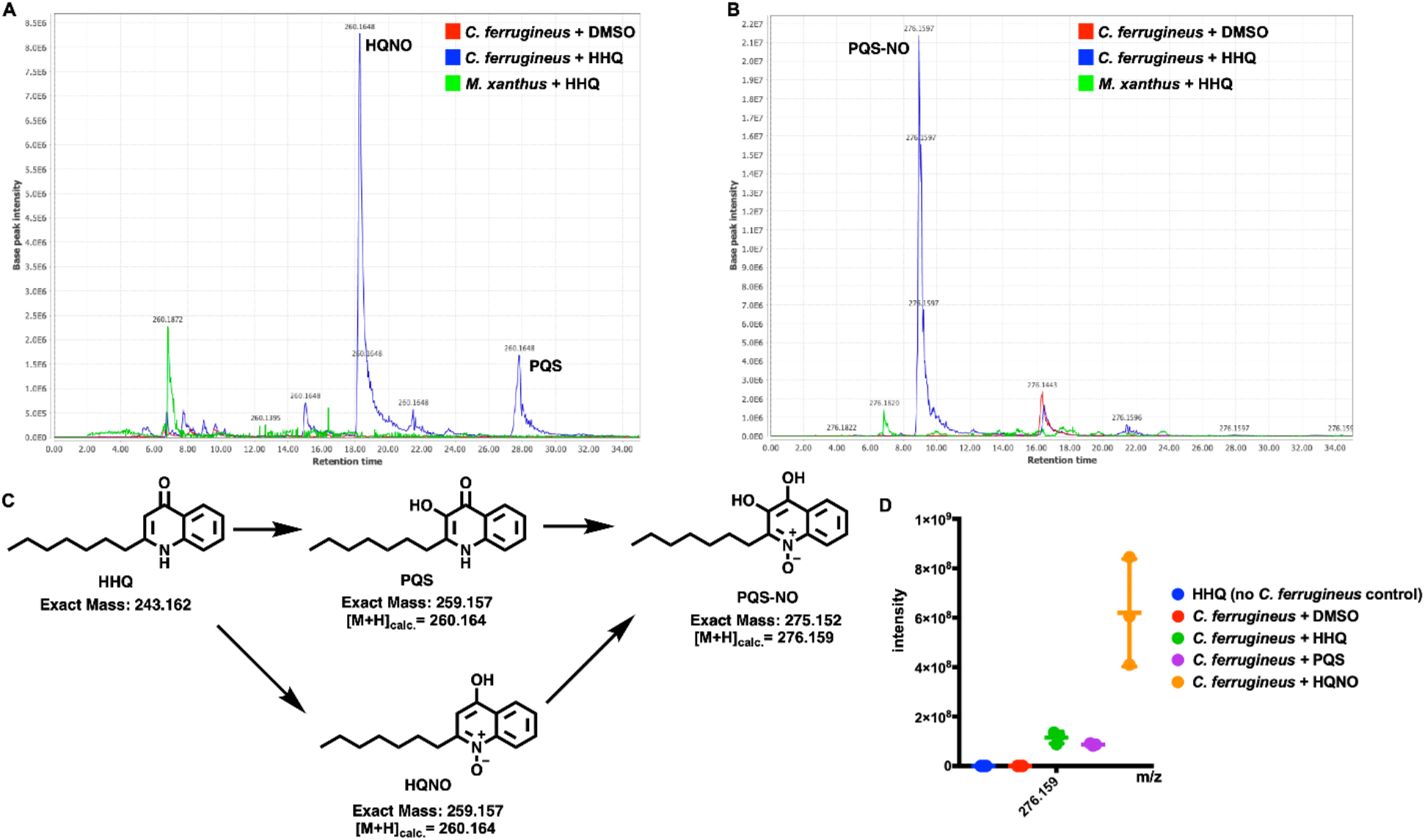
(A) Extracted ion chromatograph (EIC) depicting presence of HQNO and PQS in HHQ exposed extracts from *C. ferrugineus* and not observed in HHQ exposed extracts from *M. xanthus*. (B) EIC depicting presence of PQS-NO in HHQ exposed extracts of *C. ferrugineus*, also not present in *M. xanthus* extracts. Chromatographs rendered with MZmine v2.37. (C) Oxidative detoxification of HHQ by *C. ferrugineus* including exact mass values from ChemDraw Professional v17.1. (D) Detected ion intensities for PQS-NO comparing crude extracts of *C. ferrugineus* exposed to HHQ, PQS and HQNO; detected intensity data provided by XCMS-multigroup analysis (n=3; p ≤0.02).

### *C. ferrugineus* response to HHQ correlates with superior predation of *P. aeruginosa*

Predation assays using the lawn culture method were conducted in triplicate on lawns of *P. aeruginosa* with both *M. xanthus* and *C. ferrugineus* (Morgan et al., 2010). These assays confirmed that *P. aeruginosa* was comparatively more susceptible to predation by *C. ferrugineus* (Figure 9). These results suggest the unique response to exogenous HHQ observed from *C. ferrugineus* to be an evolved trait associated with exposure to quinolone signals that correlates with a prey range which includes quinolone signal-producing pseudomonads.

**Figure 9:**
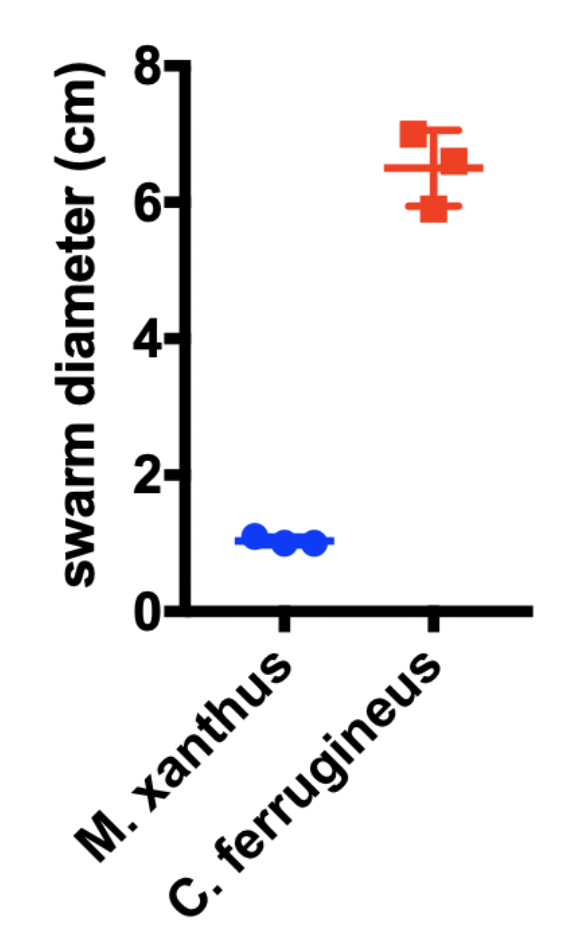
Lawn culture predation assay data depicting superior predation of *P. aeruginosa* by *C. ferrugineus* (n=3; p ≤0.005). Statistical significance calculated using an unpaired t test with Welch’s correction.

## Discussion

Although the predatory lifestyles of myxobacteria have long been associated with their capacity as a resource for natural products discovery, the chemical ecology of predator-prey interactions remains underexplored (Findlay, 2016; Munoz-Dorado et al., 2016; Herrmann et al., 2017). The recent discovery that exogenous AHL quorum signals associated with Gram-negative prey bacteria increase the predatory capacity of *M. xanthus* provides an excellent example of shared chemical space within microbial communities influencing predation (Lloyd and Whitworth, 2017). Utilizing comparative transcriptomics and metabolomics, we sought to determine the generality of predatory eavesdropping and how the phenomenon might correlate with prey range by comparing responses from *M. xanthus* and *C. ferrugineus* when exposed to structurally distinct quorum signals associated with the clinical pathogen *P. aeruginosa*.

Initial transcriptomic data comparing *M. xanthus* and *C. ferrugineus* exposed to C6-AHL revealed overlapping transcriptional responses from both myxobacteria. Originally referenced as predatory eavesdropping, we sought to determine the impact of C6-AHL on genes with annotations affiliated with predation and predatory responses such as motility features, lytic enzymes, and specialized metabolism (Munoz-Dorado et al., 2016). Transcription of multiple genes associated with transcriptional regulation, metabolism, and cell wall maintenance was influenced by exogenous C6-AHL across both myxobacteria. The only potential predatory features with a transcriptional response to C6-AHL exposure were putative lytic enzymes from *C. ferrugineus*, and no annotated genes predicted to be involved in motility were affected by C6-AHL in either myxobacteria. Considering the original observation that AHLs stimulate predation by increasing the vegetative population of *M. xanthus*, we suggest that the observed change in transcription of genes associated with metabolism and signal transduction from both myxobacteria may correspond with a similar population-based response and shift in vegetative state (Lloyd and Whitworth, 2017). The decreased transcription of the gene encoding for MrpC, a developmental regulator involved in sporulation (Robinson et al., 2014), observed in our *M. xanthus* dataset also supports a population-based response to C6-AHL.

Subsequent comparative metabolomics experiments indicated that C6-AHL and 3-oxo-C6-AHL elicit overlapping responses from both myxobacteria and that the core AHL moiety L-HSL also elicits a similar response from *C. ferrugineus*. We conclude that the overlap between C6-AHL, 3-oxo-C6-AHL, and L-HSL indicates an evolved recognition of the homoserine lactone unit present in all AHL quorum signals (Papenfort and Bassler, 2016; Mukherjee and Bassler, 2019). As generalist predators, a more general process for AHL perception that responds to a core moiety in the scaffold of AHLs might be preferred to a specialized process associated with the variable *N*-acylamides of AHLs. We also suspect this centralized response to L-HSL may relate to the absence of a LuxR-type AHL receptor that includes a conserved AHL-binding domain. The ubiquity of AHL quorum signals amongst Gram-negative bacteria combined with the overlap in observed responses from *M. xanthus* and *C. ferrugineus* suggest that AHL-based eavesdropping by myxobacteria could be a general trait that benefits predation.

Unlike the overlap in responses to AHL exposure, the quinolone signal HHQ elicited a contrasting transcriptomic from *M. xanthus* and *C. ferrugineus*. Contrary to *M. xanthus*, *C. ferrugineus* upregulated genes associated with signal transduction and transcriptional regulation, various metabolic pathways, and multiple genes associated with protein translation and turnover, cell wall biogenesis and maintenance, and specialized metabolism when exposed to HHQ. Interestingly, an annotated FAD-dependent oxidoreductase homologous to the monooxygenase PqsH from *P. aeruginosa*, which hydroxylates HHQ to yield PQS was upregulated 31-fold in *C. ferrugineus* exposed to HHQ. Subsequent metabolomic experiments confirmed the presence of two oxidized analogs of HHQ, PQS and HQNO (Dubern and Diggle, 2008; Thierbach et al., 2017). The detection of these oxidized quinolones as well as an additional feature, PQS-NO, previously associated with the oxidative detoxification of quinolone signals in HHQ exposed *C. ferrugineus* samples and absence in *M. xanthus* samples suggests that *M. xanthus* is unable to similarly metabolize HHQ (Dubern and Diggle, 2008; Thierbach et al., 2017; Ritzmann et al., 2021). Oxidative detoxification of quinolone signals produced by pseudomonads has previously been reported from strains of *Arthrobacter, Rhodococcus*, and *Staphylococcus aureus* (Thierbach et al., 2017). We suggest that this oxidative detoxification process contributes to the superior predation of *P. aeruginosa* observed from *C. ferrugineus* in our predation assays comparing both myxobacteria.

Despite being considered keystone taxa within microbial communities (Petters et al., 2021), the extent of myxobacterial bacterivory and its contribution to nutrient cycling within microbial food webs remains unknown. These results further demonstrate myxobacterial eavesdropping on prey signaling molecules and provide insight into how responses to exogenous signals might correlate with prey range of myxobacteria. Although broadly considered generalist predators, predatory specialization has been observed from myxobacteria. Oxidation of quinolone signals and superior predation of *P. aeruginosa* observed from *C. ferrugineus* provides an example of how prey signaling molecules and the shared chemical ecology of microbial communities influence myxobacterial predation.

## Experimental Procedures

### Cultivation of *M. xanthus* and *C. ferrugineus*

*Cystobacter ferrugineus* strain Cbfe23, DSM 52764, initially obtained from German Collection of Microorganisms (DSMZ) in Braunschweig, and *Myxococcus xanthus* strain GJV1 were employed in this study. *Cystobacter ferrugineus* was grown on VY/2 agar (5 g/L baker’s yeast, 1.36 g/L CaCl_2_, 0.5 mg/L vitamin B12, 15 g/L agar, pH 7.2). Whereas, CTTYE agar (1.4% w/v agar, 1% Casitone, 10 mM Tris-HCl (pH 7.6), 1 mM potassium phosphate (pH 7.6), 8 mM MgSO_4_, 0.5% yeast extract) was utilized to culture *M. xanthus*.

### Quorum signal exposure experiments

For signal exposure conditions, required volumes for 9 μM of filter sterilized, HHQ (Sigma), C6-AHL (Cayman Chemical), 3-oxo-C6-AHL (Cayman Chemical), L-HSL (Cayman Chemical), D-HSL (Cayman Chemical), PQS (Sigma), and HQNO (Sigma) from a 150 mM stock prepared in DMSO were added to autoclaved medium at 55°C. Boiled AHL samples were prepared according to established methodology (Lloyd and Whitworth, 2017). For RNA-seq and LC-MS/MS analysis, *C. ferrugineus* was cultivated on VY/2 agar medium, and *M. xanthus* was cultured on CTTYE agar medium. For all signal exposure experiments both myxobacteria were grown at 30°C with *C. ferrugineus* grown for 10 days and *M. xanthus* grown 14 days.

### RNA sequencing experiments and analysis

Myxobacterial cells were scrapped from the agar plates and stored in RNA-ladder. Total RNA was isolated from the samples using the RNeasy PowerSoil Total RNA Kit (Qiagen) following the manufacturer’s instructions. Consistent aliquots of biomass (500 mg) from each myxobacteria were used for RNA extractions. The concentration of total RNA was determined using the Qubit® RNA Assay Kit (Life Technologies). For rRNA depletion, first, 1000 ng of total RNA was used to remove the DNA contamination using Baseline-ZERO™ DNase (Epicentre) following the manufacturer’s instructions followed by purification using the RNA Clean & Concentrator-5 columns (Zymo Research). DNA free RNA samples were used for rRNA removal by using RiboMinus™ rRNA Removal Kit (Bacteria; Thermo Fisher Scientific) and final purification was performed using the RNA Clean & Concentrator-5 columns (Zymo Research). rRNA depleted samples were used for library preparation using the KAPA mRNA HyperPrep Kits (Roche) by following the manufacturer’s instructions. Following the library preparation, the final concentration of each library was measured using the Qubit® dsDNA HS Assay Kit (Life Technologies), and average library size for each was determined using the Agilent 2100 Bioanalyzer (Agilent Technologies) (Supplemental Tables 1 and 2). The libraries were then pooled in equimolar ratios of 0.75 nM, and sequenced paired end for 300 cycles using the NovaSeq 6000 system (Illumina). RNA sequencing was conducted by MR DNA (Molecular Research LP). RNAseq analysis was performed using ArrayStar V15 and the R-package DESeq2 for differential expression data. Raw data from RNAseq analysis publicly available at the National Center for Biotechnology Information Sequence Read Archive under the following BioProjects PRJNA555507, PRJNA730806, PRJNA730808.

### Organic phase extraction of metabolites

After cultivation, myxobacterial plates were manually diced and extracted with excess EtOAc. Pooled EtOAc was filtered and dried *in vacuo* to provide crude extracts for LCMS/MS analysis. LC-MS/MS analysis of the extracted samples was performed on an Orbitrap Fusion instrument (Thermo Scientific, San Jose, CA) controlled with Xcalibur version 2.0.7 and coupled to a Dionex Ultimate 3000 nanoUHPLC system. Samples were loaded onto a PepMap 100 C18 column (0.3 mm × 150 mm, 2 μm, Thermo Fisher Scientific). Separation of the samples was performed using mobile phase A (0.1% formic acid in water) and mobile phase B (0.1% formic acid in acetonitrile) at a rate of 6 μL/min. The samples were eluted with a gradient consisting of 5 to 60% solvent B over 15 min, ramped to 95% B over 2 min, held for 3 min, and then returned to 5% B over 3 min and held for 8 min. All data were acquired in positive ion mode. Collision induced dissociation (CID) was used to fragment molecules, with an isolation width of 3 m/z units. The spray voltage was set to 3600 volts, and the temperature of the heated capillary was set to 300°C. In CID mode, full MS scans were acquired from m/z 150 to 1200 followed by eight subsequent MS^2^ scans on the top eight most abundant peaks. The nominal orbitrap resolution for both the MS^1^ and MS^2^ scans was 60000. The expected mass accuracy based on external calibration was <3 ppm. MZmine 2.53 was used to generate extracted ion chromatograms (Pluskal et al., 2010).

### XCMS analysis

Generated data were converted to .mzXML files using MS-Convert (Adusumilli and Mallick, 2017). Multigroup analysis of converted .mzXML files was done using XCMS-MRM and the default HPLC/Orbitrap parameters (Domingo-Almenara et al., 2018; Forsberg et al., 2018). Within the XCMS-MRM result tables, determination of signal-impacted detected features was afforded by filtering results for those with a p ≤0.02.

#### Lawn culture predation assays

*Pseudomonas aeruginosa* ATCC 10145^T^ was purchased from the American Type Culture Collection (ATCC). The predation experiment was performed according to (Pham et al., 2005; Morgan et al., 2010). Briefly, overnight grown culture of *P. aeruginosa* was palleted at 5000 x g. The cell pellet was washed with TM buffer and pelleted again. The pelleted cells were resuspended in TM buffer to an OD600 0.5. A 250 μL volume of resuspended cell suspension was utilized to make a uniform bacterial lawn on a WAT agar plate. Myxobacterium *M. xanthus* GJV1 was grown on CTTYE agar, and *C. ferrugineus* was grown on VY/2 agar for 7 days. A 600 mm^2^ agar block of each myxobacteria was excised and placed at the center of the *P. aeruginosa* cell lawn. Assays were incubated at 30°C and swarm diameters measured after 4 days.

## Acknowledgements

The authors appreciate funding and support from the National Institute of Allergy and Infectious Diseases (R15AI137996). Research reported in this publication was supported by an Institutional Development Award (IDeA) from the National Institute of General Medical Sciences of the National Institutes of Health under award number P20GM130460. The authors also appreciate Dr. Peter Zee for providing us with *M. xanthus* strain GJV1.

