## Supplemental Material for "Differential response to prey quorum signals indicates predatory range of myxobacteria"

**Supplemental Table 1.** Concentration of RNA, final library concentration, and average library size for *C. ferrugineus* RNAseq samples

| Sample | RNA Concentration (ng/uL) | Library Concentration (ng/uL) | Avg Library Size (bp) |
| --- | --- | --- | --- |
| CystobacterferrugineusDMSO1 | 864.0 | 36.00 | 427 |
| CystobacterferrugineusDMSO2 | 1016.0 | 37.60 | 411 |
| CystobacterferrugineusDMSO3 | 896.0 | 41.60 | 418 |
| CystobacterferrugineusHHQ1 | 832.0 | 38.40 | 404 |
| CystobacterferrugineusHHQ2 | 744.0 | 41.80 | 426 |
| CystobacterferrugineusHHQ3 | 804.0 | 49.00 | 450 |
| CystobacterferrugineusC6HSL1 | 904.00 | 48.00 | 550 |
| CystobacterferrugineusC6HSL2 | 1060.00 | 33.20 | 477 |
| CystobacterferrugineusC6HSL3 | 960.00 | 55.00 | 495 |

**Supplemental Table 2.** Concentration of RNA, final library concentration, and average library size for *M. xanthus* RNAseq samples

| Sample | RNA Concentration (ng/uL) | Library Concentration (ng/uL) | Avg Library Size (bp) |
| --- | --- | --- | --- |
| M-xanthus-GJV1-DMSO-1 | 86.4 | 51.20 | 462 |
| M-xanthus-GJV1-DMSO-2 | 66.4 | 50.80 | 470 |
| M-xanthus-GJV1-DMSO-3 | 120.0 | 51.60 | 476 |
| M-xanthus-GJV1-HHQ-1 | 72.6 | 46.00 | 461 |
| M-xanthus-GJV1-HHQ-2 | 112.0 | 55.00 | 470 |
| M-xanthus-GJV1-HHQ-3 | 200.0 | 47.00 | 476 |
| M-xanthus-GJV1-C6HSL-1 | 48.4 | 53.00 | 468 |
| M-xanthus-GJV1-C6HSL-2 | 430.0 | 56.40 | 479 |
| M-xanthus-GJV1-C6HSL-3 | 430.0 | 53.00 | 505 |

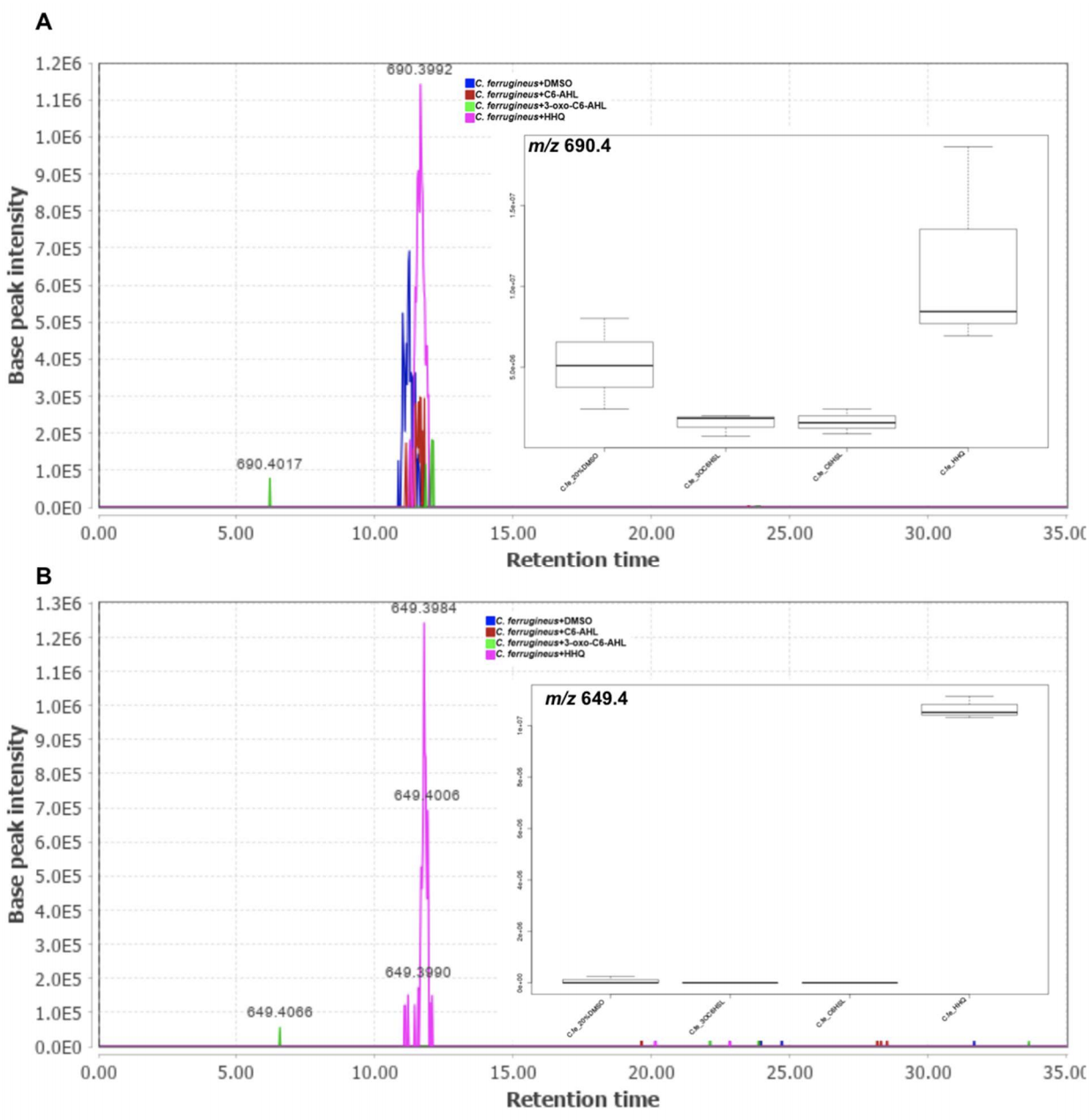
**Supplemental Figure 1:** Exemplary box plots of XCMS data used to generate Venn diagrams in Figure 5. (A) An extracted ion chromatograph (EIC) of one of the impacted metabolic features (m/z 690.4) with its corresponding box-plot from XCMS statistical analysis. *C. ferrugineus* crude extracts depicting increased detected ion intensity for the feature detected at 690.4 m/z when exposed to HHQ and decreased detected ion intensity when exposed to AHL signals compared to signal unexposed (DMSO) control sample. (B) An EIC of one of the impacted metabolic features (m/z 649.4), with its corresponding box-plot from XCMS statistical analysis, detected exclusively in HHQ exposed *C. ferrugineus* samples. Chromatograph rendered with MZmine v2.37. Box-plot data is provided by XCMS-multigroup analysis (n=3, p ≤0.02). On the bar-graph, the x-axis shows different exposure conditions, and y-axis shows base peak intensity of a detected ion.

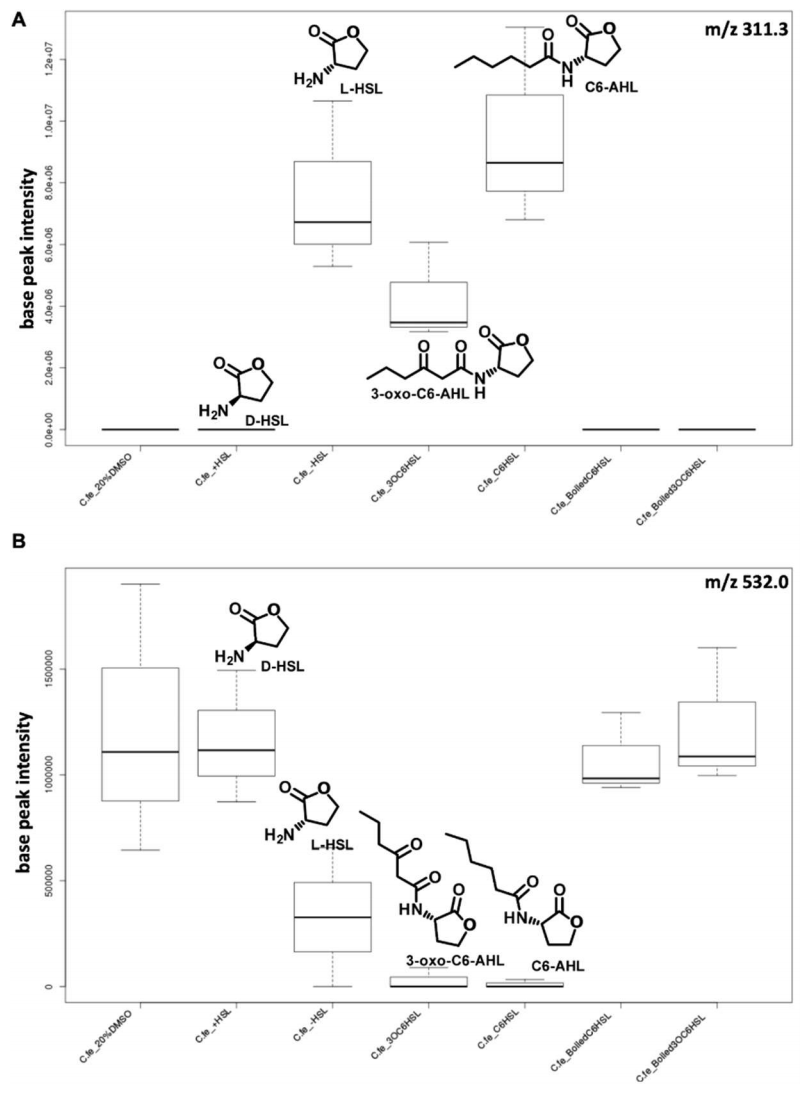

**Supplemental Figure 2:** Box-plot data from XCMS showing *C. ferrugineus* overlapping response to L-homoserine lactone and acylhomoserine lactones. (A, B) Examples of two of the impacted metabolic features (311.3 m/z & 532.0 m/z) impacted similarly by L-HSL, C-6-AHL, and 3-oxo-C6-AHL whereas ion intensities for the same features in case of D-HSL, and boiled AHLs (2 right box-plots) exposure correspond to signal unexposed (DMSO) controls. Box-plot data is provided by XCMS-multigroup analysis (n=3, p ≤0.02).

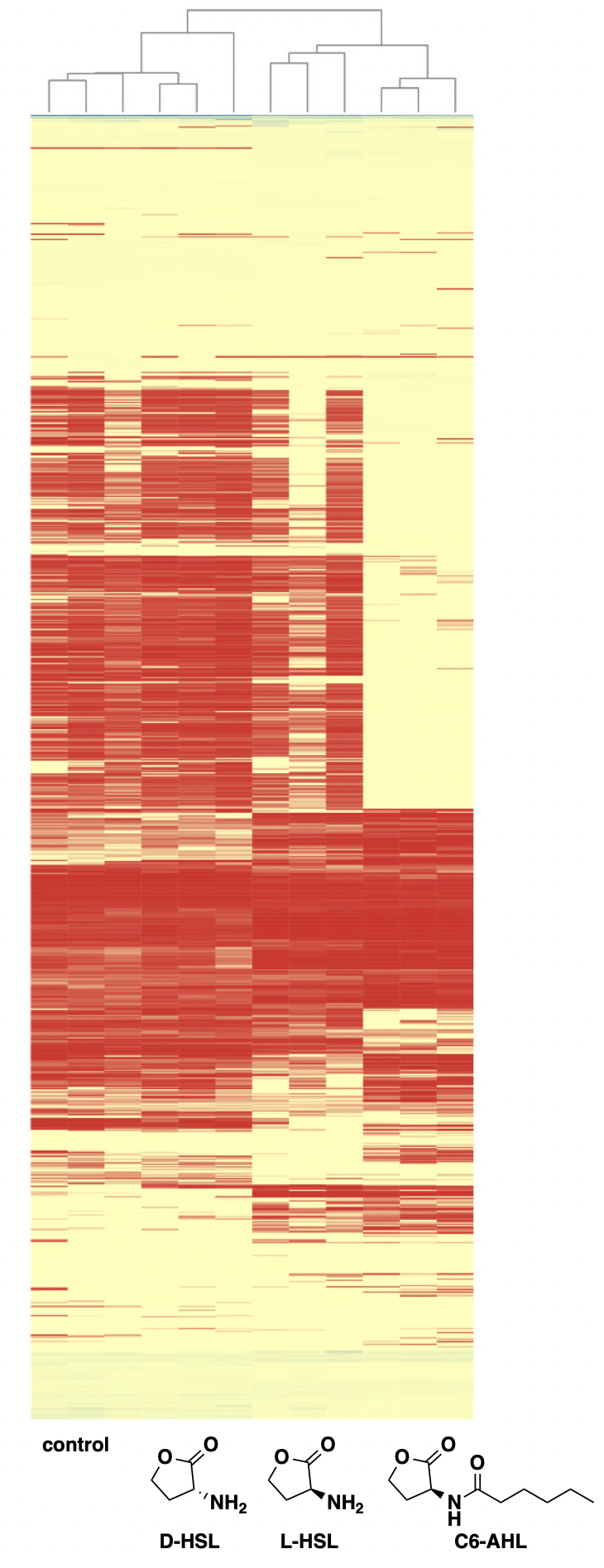

**Supplemental Figure 3:** Hierarchical clustering of a detected feature intensity heat map rendered in XCMS depicting clustering of L-HSL response with C6-AHL response observed from *C. ferrugineus*.

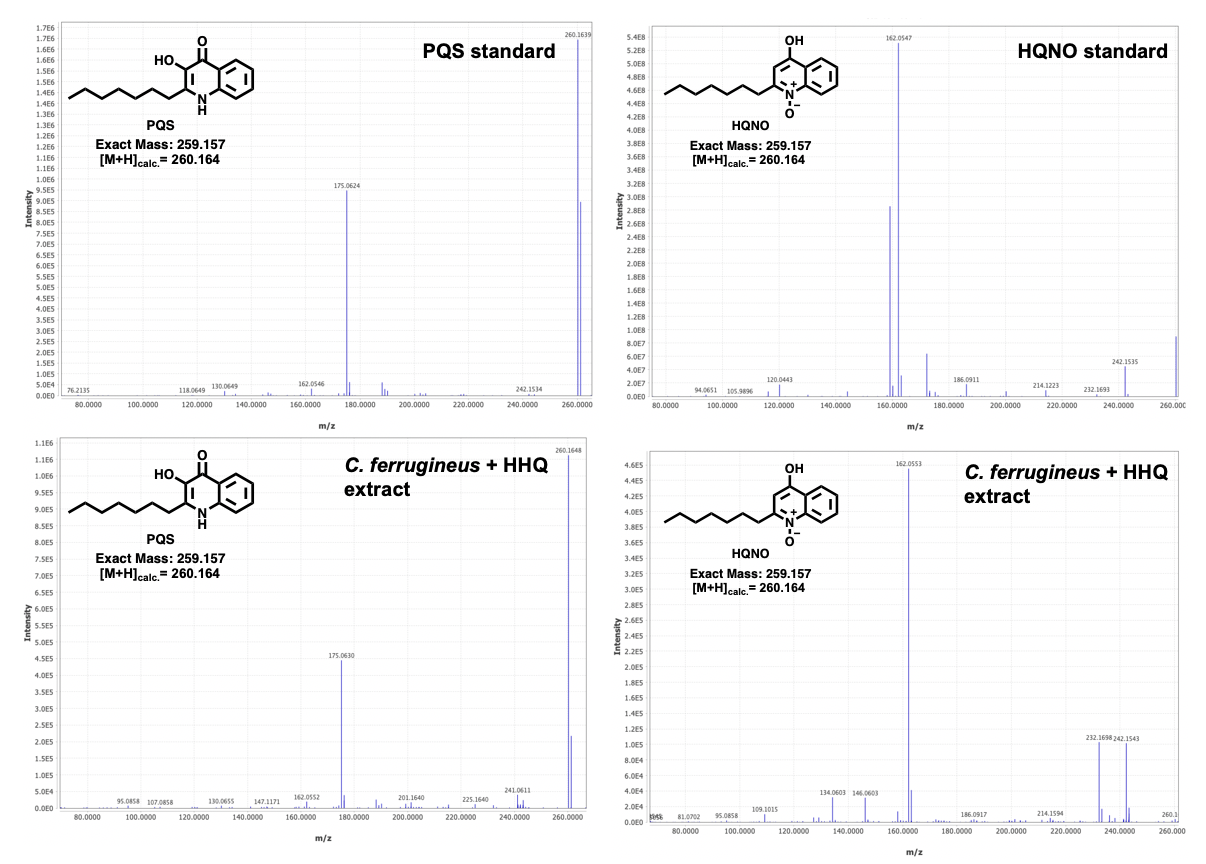
**Supplemental Figure 4:** Comparison of MS^2^ fragmentation patterns from LC-MS/MS analysis PQS and HQNO commercial standards and PQS and HQNO detected in HHQ exposed *C. ferrugineus* extracts.

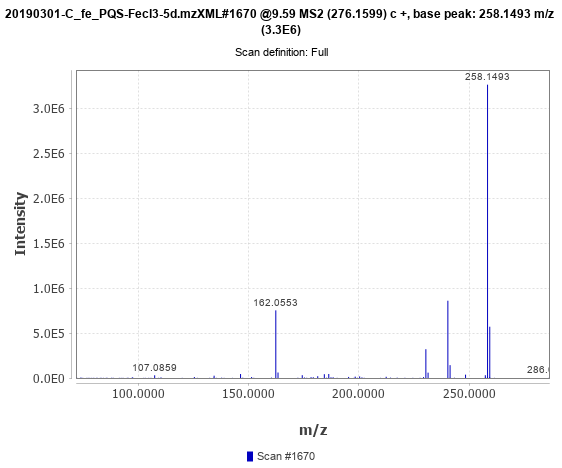

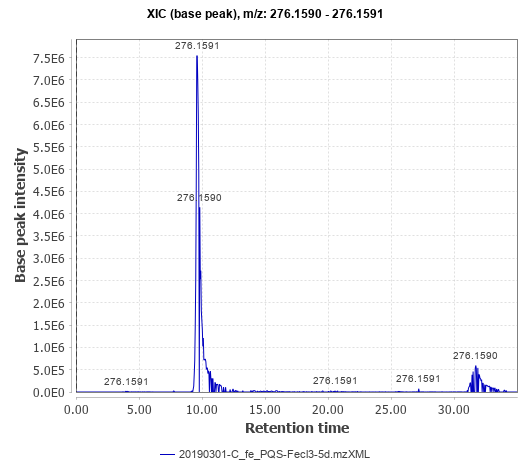

**Supplemental Figure 5:** EIC of putative PQS-NO product and corresponding MS^2^ fragmentation pattern from LC-MS/MS analysis of PQS exposed *C. ferrugineus* extracts.

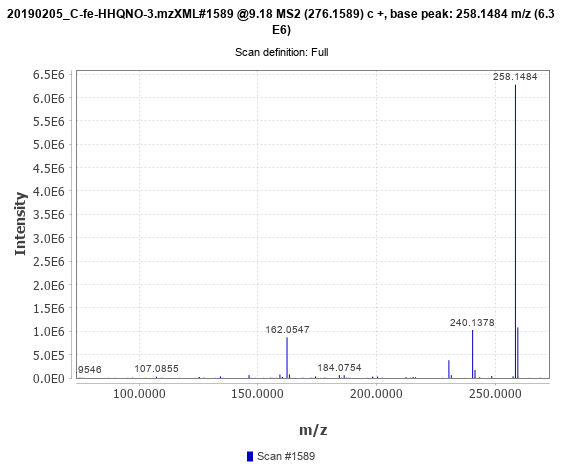

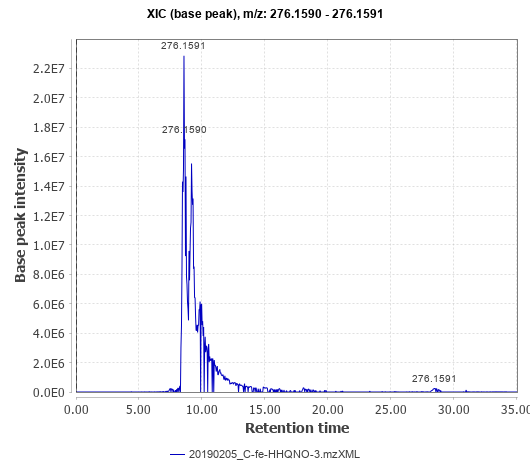

**Supplemental Figure 6:** EIC of putative PQS-NO product and corresponding MS^2^ fragmentation pattern from LC-MS/MS analysis of HQNO exposed *C. ferrugineus* extracts.
